# High-throughput annotation of full-length long noncoding RNAs with Capture Long-Read Sequencing

**DOI:** 10.1101/105064

**Authors:** Julien Lagarde, Barbara Uszczynska-Ratajczak, Silvia Carbonell, SÍlvia Pérez-Lluch, Amaya Abad, Carrie Davis, Thomas R. Gingeras, Adam Frankish, Jennifer Harrow, Roderic Guigo, Rory Johnson

**Affiliations:** Centre for Genomic Regulation (CRG), The Barcelona Institute of Science and Technology, Dr. Aiguader 88, 08003 Barcelona, Spain.; Universitat Pompeu Fabra (UPF), Barcelona, Spain.; R&D Department, Quantitative Genomic Medicine Laboratories (qGenomics), Barcelona, Spain.; Functional Genomics Group, Cold Spring Harbor Laboratory, 1 Bungtown Road, Cold Spring Harbor, New York 11724, USA.; Wellcome Trust Sanger Institute, Hinxton, Cambridgeshire, UK CB10 1HH.; Present address: Centre of New Technologies, S. Banacha 2C, 02-097 Warsaw, Poland; Present address: Illumina, Cambridge, UK.; Present address: Department of Clinical Research, University of Bern, Murtenstrasse 35, 3010 Bern, Switzerland.

**Keywords:** Long noncoding RNA, lncRNA, lincRNA, RNA sequencing, transcriptomics, GENCODE, annotation, CaptureSeq, third generation sequencing, long read sequencing, PacBio, KANTR

## Abstract

Accurate annotations of genes and their transcripts is a foundation of genomics, but no annotation technique presently combines throughput and accuracy. As a result, reference gene collections remain incomplete: many gene models are fragmentary, while thousands more remain uncatalogued–particularly for long noncoding RNAs (lncRNAs). To accelerate lncRNA annotation, the GENCODE consortium has developed RNA Capture Long Seq (CLS), combining targeted RNA capture with third-generation long-read sequencing. We present an experimental re-annotation of the GENCODE intergenic lncRNA population in matched human and mouse tissues, resulting in novel transcript models for 3574 / 561 gene loci, respectively. CLS approximately doubles the annotated complexity of targeted loci, outperforming existing short-read techniques. Full-length transcript models produced by CLS enable us to definitively characterize the genomic features of lncRNAs, including promoter- and gene-structure, and protein-coding potential. Thus CLS removes a longstanding bottleneck of transcriptome annotation, generating manual-quality full-length transcript models at high-throughput scales.

**Abbreviations:**

bp: base pair
FL: full length
nt: nucleotide
ROI: read of insert, *i.e.* PacBio read
SJ: splice junction
SMRT: single-molecule real-time
TM: transcript model

## Introduction

Long noncoding RNAs (lncRNAs) represent a vast and largely unexplored component of the mammalian genome. Efforts to assign lncRNA functions rest on the availability of high-quality transcriptome annotations. At present such annotations are still rudimentary: we have little idea of the total lncRNA count, and for those that have been identified, transcript structures remain largely incomplete.

The number and size of available lncRNA annotations have grown rapidly thanks to projects using diverse approaches. Early gene sets, deriving from a mixture of FANTOM cDNA sequencing efforts and public databases^1,2^ were joined by the “lincRNA” (long intergenic non-coding RNA) sets, discovered through chromatin signatures^3^. More recently, studies have applied transcript-reconstruction software, such as *Cufflinks*^4^ to identify novel genes in short-read RNA sequencing (RNAseq) datasets^5–9^. However the reference for lncRNAs has become the regularly-updated, manual annotations from GENCODE, based on curation of cDNAs/ESTs by human annotators^10,11^, and adopted by international genomics consortia^12–15^.

At present, annotation efforts are caught in a trade-off between throughput and quality. Short read-based transcriptome reconstruction methods deliver large annotations with low financial and time investment. Manual annotation is slow and requires longterm funding. However the quality of software-reconstructed annotations is often doubtful, due to the inherent difficulty of reconstructing transcript structures from shorter sequence reads. Such structures tend to be incomplete, often lacking terminal exons or omitting splice junctions between adjacent exons^16^. This particularly affects lncRNAs, whose low expression results in low read coverage^11^. The outcome is a growing divergence between large automated annotations of uncertain quality (*e.g*. 101,700 genes for NONCODE^8^), and the highly-curated “conservative” GENCODE collection^11^ (15,767 genes for version 25).

Annotation incompleteness takes two forms. First, genes may be entirely missing from the annotation: many genomic regions are suspected to transcribe RNA but contain no annotation, including “orphan” small RNAs with presumed long precursors^17^, enhancers^18^ and ultraconserved elements^19,20^. Second, annotated lncRNAs may represent partial gene structures. Start and end sites frequently lack independent supporting evidence^11^, and lncRNAs are shorter and have fewer exons than mRNAs^7,11,21^. Recently, RACE-Seq was developed to complete lncRNA annotations, but at relatively low throughput^21^.

One of the principal impediments to lncRNA annotation arises from their low steady-state levels^3,11^. To overcome this, “RNA Capture Sequencing” (CaptureSeq)^22^ is used to boost the concentration of low-abundance transcripts in cDNA libraries. These studies depend on short-read sequencing and *in-silico* transcript reconstruction^22–24^. Thus, while CaptureSeq achieves high throughput, its transcript structures lack the confidence required for inclusion in GENCODE.

In this study, we describe a new method called RNA Capture Long Seq (CLS), which couples targeted RNA capture with third-generation long-read cDNA sequencing. We use CLS to interrogate the GENCODE catalogue of intergenic lncRNAs, together with thousands of suspected novel loci, in six tissues each of human and mouse. CLS couples the throughput of CaptureSeq with high-confidence, complete transcript models from long-read sequencing, resulting in a significant advance in transcriptome annotation.

## Results

### Capture Long Seq approach to complete lncRNA annotations

Our aim was to develop an experimental approach to improve and extend reference transcript annotations, while minimizing human intervention and avoiding *in-silico* transcript assembly. We designed a method, Capture Long Seq (CLS), which couples targeted RNA capture to Pacific Biosciences (“PacBio”) Third Generation long-read sequencing (Figure 1a).

**Figure 1:**
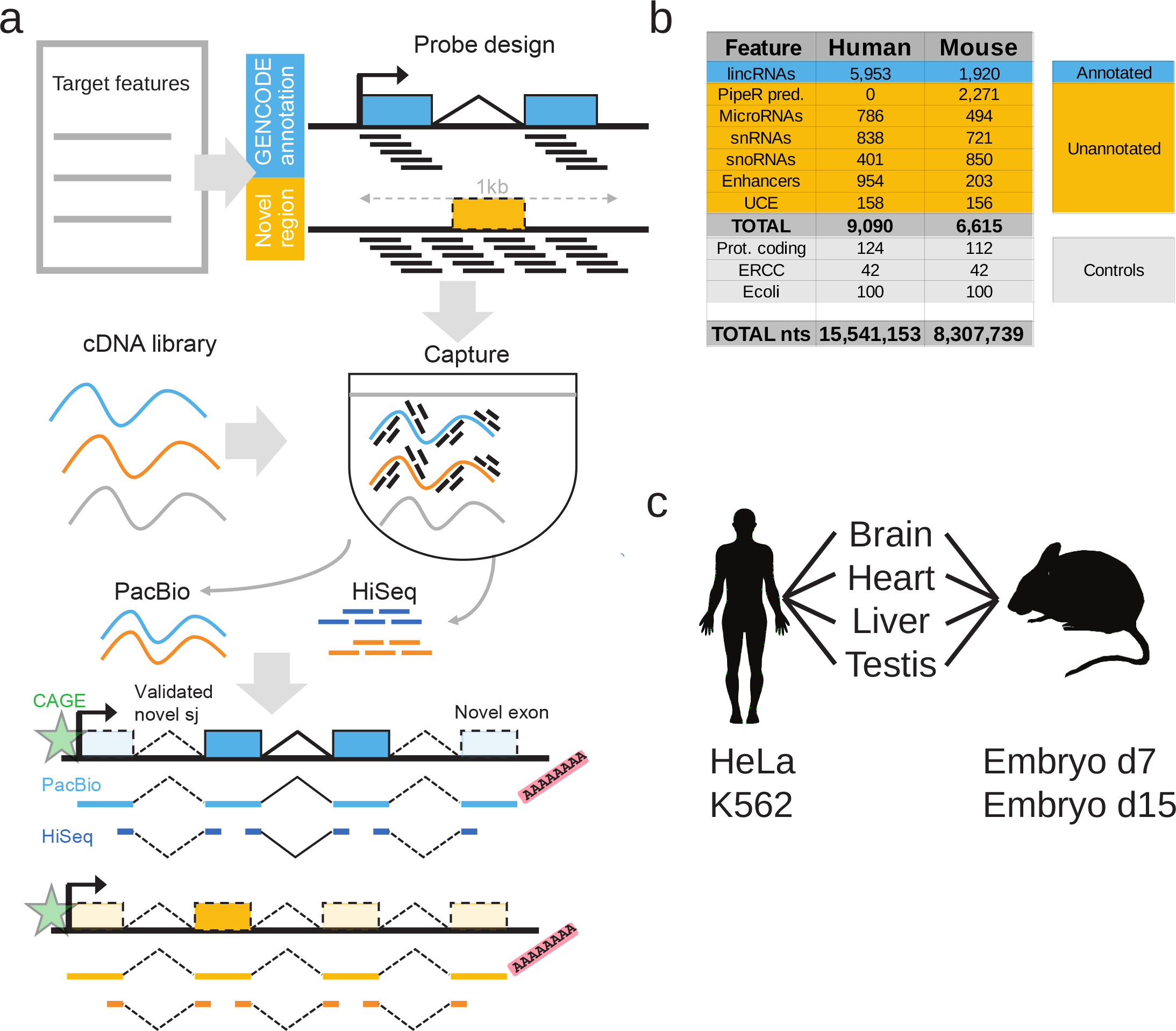
Capture Long Seq approach to extend the GENCODE lncRNA annotation. **a)** Strategy for automated, high-quality transcriptome annotation. CLS may be used to complete existing annotations (blue), or to map novel transcript structures in suspected loci (orange). Capture oligonucleotides (black bars) are designed to tile across targeted regions. PacBio libraries are prepared for from the captured molecules. Illumina HiSeq short-read sequencing can be performed for independent validation of predicted splice junctions. Predicted transcription start sites can be confirmed by CAGE clusters (green), and transcription termination sites by non-genomically encoded polyA sequences in PacBio reads. Novel exons are denoted by lighter coloured rectangles. **b)** Summary of human and mouse capture library designs. Shown are the number of individual gene loci that were probed. “PipeR pred.”: orthologue predictions in mouse genome of human lncRNAs, made by PipeR^29^; “UCE”: ultraconserved elements; “Prot. coding”: expression-matched, randomly-selected protein-coding genes; “ERCC”: spike-in sequences; “Ecoli”: randomly-selected *E. coli* genomic regions. Enhancers and UCEs are probed on both strands, and these are counted separately. “Total nts”: sum of targeted nucleotides. **c)** RNA samples used.

CLS may be applied to two distinct objectives: to improve existing gene models, or to identify novel loci (blue and orange in Figure 1a, respectively). Although the present study focuses mainly on the former, we demonstrate also that novel loci can be captured and sequenced. We created a comprehensive capture library targeting the set of intergenic GENCODE lncRNAs in human and mouse. Note that annotations for human are presently more complete than for mouse, resulting in different annotation sizes (14,470 vs 5,385 lncRNA genes in GENCODE releases 20 and M3, respectively). GENCODE annotations probed in this study are principally multi-exonic transcripts based on polyA+ cDNA/EST libraries, and hence not likely to include “enhancer RNAs”^10,25^. To these we added tiled probes targeting loci that may produce lncRNAs: small RNA genes^26^, enhancers^27^ and ultraconserved elements^28^. For mouse we also added orthologous lncRNA predictions from PipeR^29^. Numerous control probes were added, including a series targeting half of the ERCC synthetic spike-ins^30^. These sequences were targeted by capture libraries of temperature-matched and non-repetitive oligonucleotide probes (Figure 1b).

To access the maximal lncRNA diversity, we chose transcriptionally complex and biomedically-relevant organs from mouse and human: whole brain, heart, liver and testis (Figure 1c). We added two deeply-studied human cell lines, HeLa and K562^31^, and two mouse embryonic time-points (E7 and E15).

We designed a protocol to capture full-length, oligo-dT-primed cDNAs (see Methods). Barcoded, unfragmented cDNAs were pooled and captured. Preliminary tests using qPCR indicated enrichment for targeted regions (Supplementary Figure 1a). PacBio sequencing tends to favour shorter templates in a mixture^32^. Therefore pooled, captured cDNA was size-selected into three ranges (1-1.5kb, 1.5-2.5kb, >2.5kb) (Supplementary Figure 1b-c), and used to construct sequencing libraries for PacBio SMRT (single-molecular real-time) technology^33^.

### CLS yields an enriched long-read transcriptome

Samples were sequenced on 130 SMRT cells, yielding ~2 million reads in total in each species (Figure 2a). PacBio reads, or “reads of insert” (ROIs) were demultiplexed to retrieve their tissue of origin and mapped to the genome. We observed high mapping rates (>99% in both cases), of which 86% and 88% were unique, in human and mouse, respectively (Supplementary Figure 2a). For brevity, all data are henceforth quoted in order of human then mouse. The use of short barcodes meant that, for ~30% of reads, the tissue of origin could not be retrieved (Supplementary Figure 2b). This may be remedied by the use of longer barcodes. Representation was even across tissues, with the exception of testis (Supplementary Figure 2d). ROIs had a median length of 1 - 1.5 kb (Figure 2b) consistent with previous reports^32^ and exceeding average lncRNA annotation of ~0.5 kb^11^.

**Figure 2:**
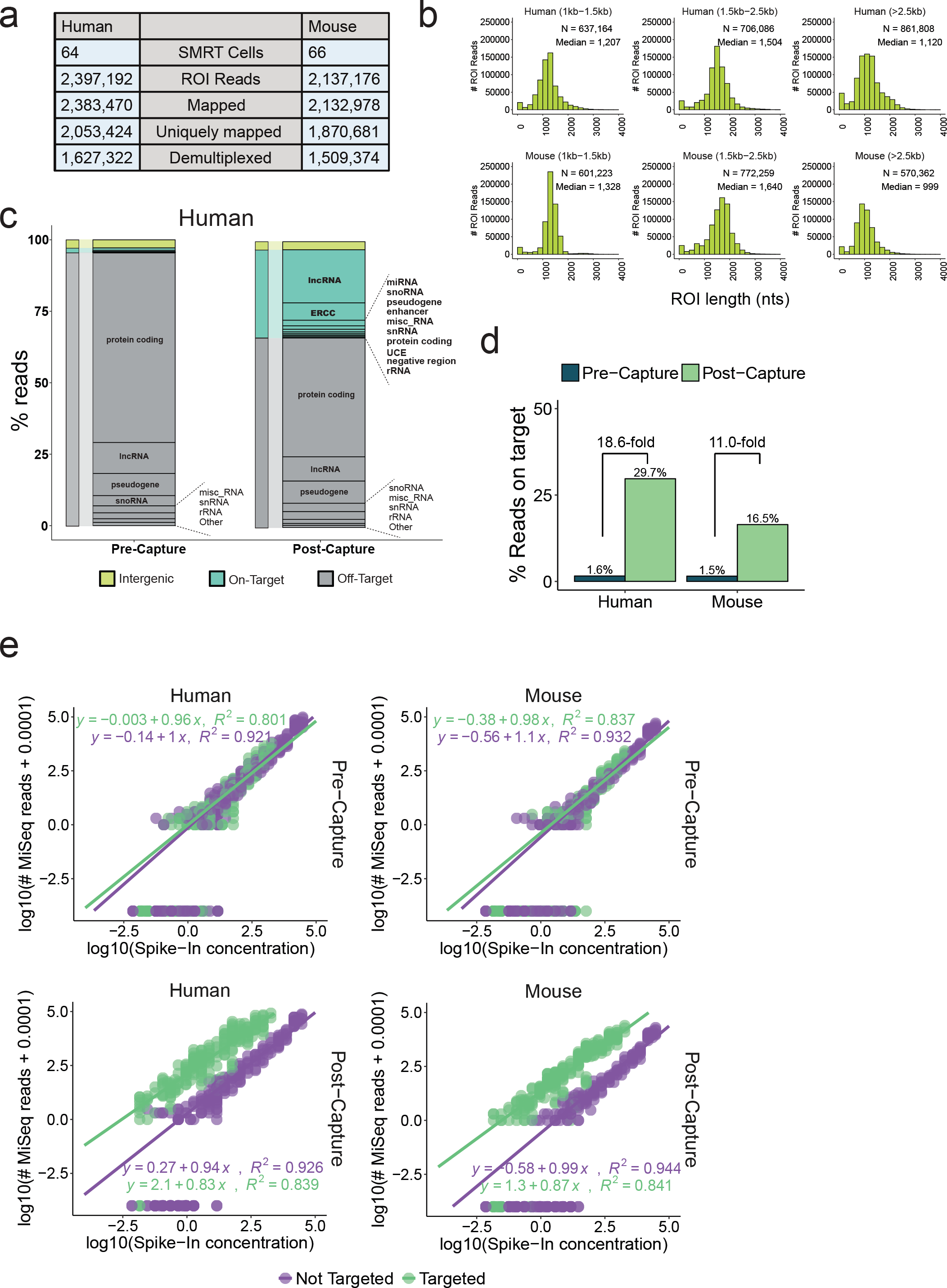
CLS yields an enriched, long-read transcriptome. **a)** Sequencing statistics. ROI = “Read Of Insert”, or PacBio reads. **b)** Length distributions of ROIs. Sequencing libraries were prepared from three size-selected cDNA fractions (see Supplementary Figure 1b-c). **c)** Breakdown of sequenced reads by gene biotype, pre- (left) and post-capture (right), for human (equivalent mouse data in Supplementary Figure 2j). Colours denote the on/off-target status of the reads: Green: reads from targeted features, including lncRNAs; Grey: reads originating from annotated but not targeted features; Yellow: reads from unannotated, non-targeted regions. The ERCC class comprises only those ERCC spike-ins that were probed. Note that when a given read overlapped more than one targeted class of regions, it was counted in each of these classes separately. **d)** Summary of capture performance. The y-axis shows percent of all mapped ROIs originating from a targeted region (“on-target”). Enrichment is defined as the ratio of this value in Post- and Pre-capture samples. Sequencing was performed using MiSeq technology. **e)** Response of read counts in captured cDNA to input RNA concentration. Upper panels: Pre-capture; Lower panels: Post-capture. Left: human; right: mouse. Note log scales for each axis. Points represent 92 spiked-in synthetic ERCC RNA sequences. 42 were probed in the capture design (green), the other 50 were not (violet). Lines represent linear fits to each dataset, whose parameters are shown. Given the log-log representation, a linear response of read counts to template concentrate should yield an equation of type *y* = *c* + *mx*, where *m* is 1.

Capture performance is assessed in two ways: by “on-target” rate – the proportion of reads originating from probed regions – and by enrichment, or increase of on-target rate following capture^34^. To estimate these, we sequenced pre- and post-capture libraries using MiSeq. CLS achieved on-target rates of 29.7% / 16.5%, representing 19- / 11-fold enrichment (Figure 2c, d and Supplementary Figure 2e). These rates are competitive with intergenic lncRNA capture in previous, short-read studies (Supplementary Figure 2f-g). The majority of off-target signal arises from non-targeted, annotated protein-coding genes (Figure 2c).

CLS on-target rates were comparable to previous studies using fragmented cDNA^35^(Supplementary Figure 2f-g), although lower than genomic DNA capture. Side-by-side comparisons showed that capturing long cDNA fragments implies some loss in capture efficiency (Supplementary Figure 2h-i), as observed by others^24^.

Synthetic spike-in sequences at known concentrations were used to assess sensitivity and quantitativeness. We compared the relationship of sequence reads to starting concentration for the 42 probed (green) and 50 non-probed (violet) synthetic ERCC sequences in pre- and post-capture samples (Figure 2e, top and bottom rows). CLS is surprisingly sensitive, extending detection sensitivity by two orders of magnitude, and capable of detecting molecules at approximately 5 × 10^−3^ copies per cell (Methods). It is less quantitative than CaptureSeq^24^, particularly at higher concentrations where the slope falls below unity. This suggests saturation of probes by cDNA molecules during hybridisation. A degree of noise, as inferred by the coefficient of determination (R^2^) between read counts and template concentration, is introduced by the capture process.

### CLS expands the complexity of known and novel lncRNAs

CLS discovers a wealth of novel transcript structures within annotated lncRNA loci. In the *SAMMSON* oncogene^36^ (*LINC01212*), we discover new exons, splice sites, and transcription termination sites compared to present annotations (Figure 3a, more examples in Supplementary Figures 3, 4, 5, which could be validated by RT-PCR).

**Figure 3:**
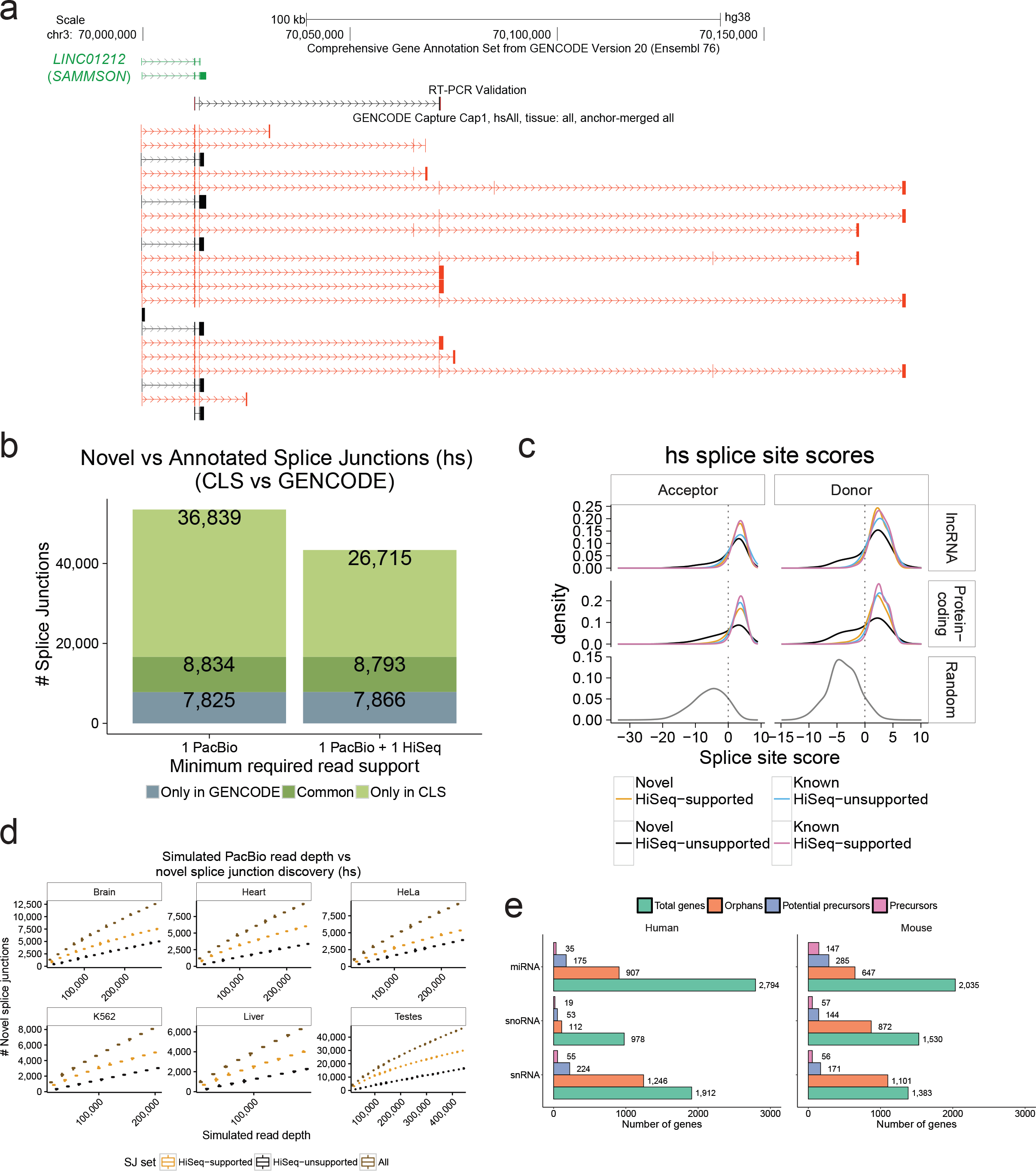
Extending known lncRNA gene structures. **a)** Novel transcript structures from the *SAMMSON* locus. Green: GENCODE; Black/Red: known/novel CLS transcript models (TMs), respectively. An RT-PCR-amplified sequence is shown. **b)** Splice junction (SJ) discovery. *Y*-axis: unique SJs for human (mouse data in Supplementary Figure 6b) within probed lncRNA loci. Grey: GENCODE-annotated, CLS-undetected SJs. Dark green: CLS-detected, GENCODE-annotated SJs. Light green: novel CLS SJs. Left: all SJs; Right: high-confidence, HiSeq-supported SJs. See Supplementary Figure 6c for comparison to the *miTranscriptome* catalogue. **c)** Splice junction (SJ) motif strength. Panels plot the distribution of predicted SJ strength, for splice site (SS) acceptors (left) and donors (right) in human (mouse data in Supplementary Figure 7a). SS strength was computed using GeneID^37^. Data are shown for non-redundant CLS SJs from targeted lncRNAs (top), protein-coding genes (middle), or randomly-selected SS-like dinucleotides (bottom). **d)** Splice junction discovery/saturation analysis in human. Panels show novel SJs discovered (*y*-axis) in simulations with increasing numbers of randomly sampled CLS ROIs (*x*-axis). SJs retrieved in each sample were stratified by level of support (Brown: all PacBio SJs; Orange: HiSeq-supported; Black: HiSeq-unsupported). Boxplots summarise 50 samples. Equivalent mouse data in Supplementary Figure 8a, and for novel TM discovery in Supplementary Figure 8b. **e)** Identification of putative precursor transcripts of small RNA genes. For each gene biotype, figures show the count of unique genes. “Orphans”: no annotated overlapping transcript in GENCODE, and targeted in capture library. “Potential Precursors”: orphan RNAs residing in the intron of a novel CLS TM. “Precursors”: reside in the exon of a novel transcript.

We measured the amount of new complexity discovered in targeted lncRNA loci. CLS detected 58% and 45% of targeted lncRNA nucleotides, and extended these annotations by 6.3 / 1.6 Mb nucleotides (86% / 64% increase compared to existing annotations) (Supplementary Figure 6a). CLS discovered 45,673 and 11,038 distinct splice junctions (SJs), of which 36,839 and 8,847 are novel (Figure 3b and Supplementary Figure 6b, left bars). The number of novel, high-confidence SJs amounted to 20,327 when compared to a deeper human SJ reference catalogue composed of both GENCODE v20 and miTranscriptome^7^ (Supplementary Figure 6c). For independent validation, and given the relatively high sequence indel rate detected in PacBio reads (Supplementary Figure 2m) (see Methods for analysis of sequencing error rates), we deeply sequenced captured cDNA by Illumina HiSeq at an average depth of 35 million / 26 million pair-end reads per sample. Split reads from this data exactly matched 78% / 75% SJs from CLS. These “high-confidence” SJs alone represent a 160% / 111% increase over the existing, probed annotations (Figure 3b, Supplementary Figure 6b). Novel high-confidence lncRNA SJs are rather tissue-specific, with greatest numbers observed in testis (Supplementary Figure 6d), and were also discovered across other classes of targeted and non-targeted loci (Supplementary Figure 6e). We observed a greater frequency of intron retention events in lncRNAs, compared to protein-coding transcripts (Supplementary Figure 6f).

To evaluate the biological significance of novel lncRNA SJs, we computed their strength using standard position weight matrix models^37^ (Figure 3c, Supplementary Figure 7a). High-confidence novel SJs from lncRNAs (orange, upper panel) far exceed the predicted strength of background SJ-like dinucleotides (bottom panels), and are essentially indistinguishable from annotated SJs (pink, upper and middle panels). Even unsupported, novel SJs (black) tend to have high scores, although with a significant low-scoring tail. Although they display little evidence of sequence conservation using standard measures (similar to lncRNA SJs in general) (Supplementary Figure 7b), novel SJs also display weak but non-random evidence of selected function (Supplementary Figure 7c).

We estimated how close these sequencing data are to saturation, *i.e*., to reaching a definitive annotation. We tested the rate of novel splice junction and transcript model discovery as a function of increasing depth of randomly-sampled ROIs (Figure 3d, Supplementary Figures 8a-b). We observed an ongoing gain of novelty with increasing depth, for both low- and high-confidence SJs, up to that presented here. Similarly, no SJ discovery saturation plateau was reached at increasing simulated HiSeq read depth (Supplementary Figure 8c). Thus, considerable additional sequencing is required to complete existing lncRNA gene structures.

Beyond lncRNAs, CLS can be used to characterize other types of transcriptional units. As an illustration, we searched for precursors of small RNAs, whose annotation remains poor^17^. We probed 1 kb windows around all “orphan” small RNAs, *i.e*. those with no annotated overlapping transcript. Note that, although mature snoRNAs are non-polyadenylated, they are processed from polyA+ precursors^38^. We identified more than one hundred likely primary transcripts, and hundreds more potential precursors harbouring small RNAs within their introns (Figure 3e). One intriguing example was the cardiac-enriched hsa-mir-143, for which CLS identifies a new RT-PCR-supported primary transcript that belongs to the *CARMEN1* lncRNA gene^39^ (Supplementary Figure 9).

### Assembling a full-length lncRNA annotation

A unique benefit of the CLS approach is the ability to identify full-length transcript models with confident 5’ and 3’ termini. ROIs of oligo-dT-primed cDNAs carry a fragment of the poly(A) tail, which can identify the polyadenylation site with basepair precision^32^. Using conservative filters, 73% / 64% of ROIs had identifiable polyA sites (Supplementary Table 1) representing 16,961 / 12,894 novel sites when compared to end positions of GENCODE annotations. Known and novel polyA sites were preceded by canonical polyadenylation motifs (Supplementary Figure 10a-d). Similarly, the 5’ completeness of ROIs was confirmed by proximity to methyl-guanosine caps identified by CAGE (Cap Analysis of Gene Expression)^15^ (Supplementary Figure 10e). CAGE and polyA sites were used to define the 5’ / 3’ completeness of all ROIs (Figure 4a).

**Figure 4:**
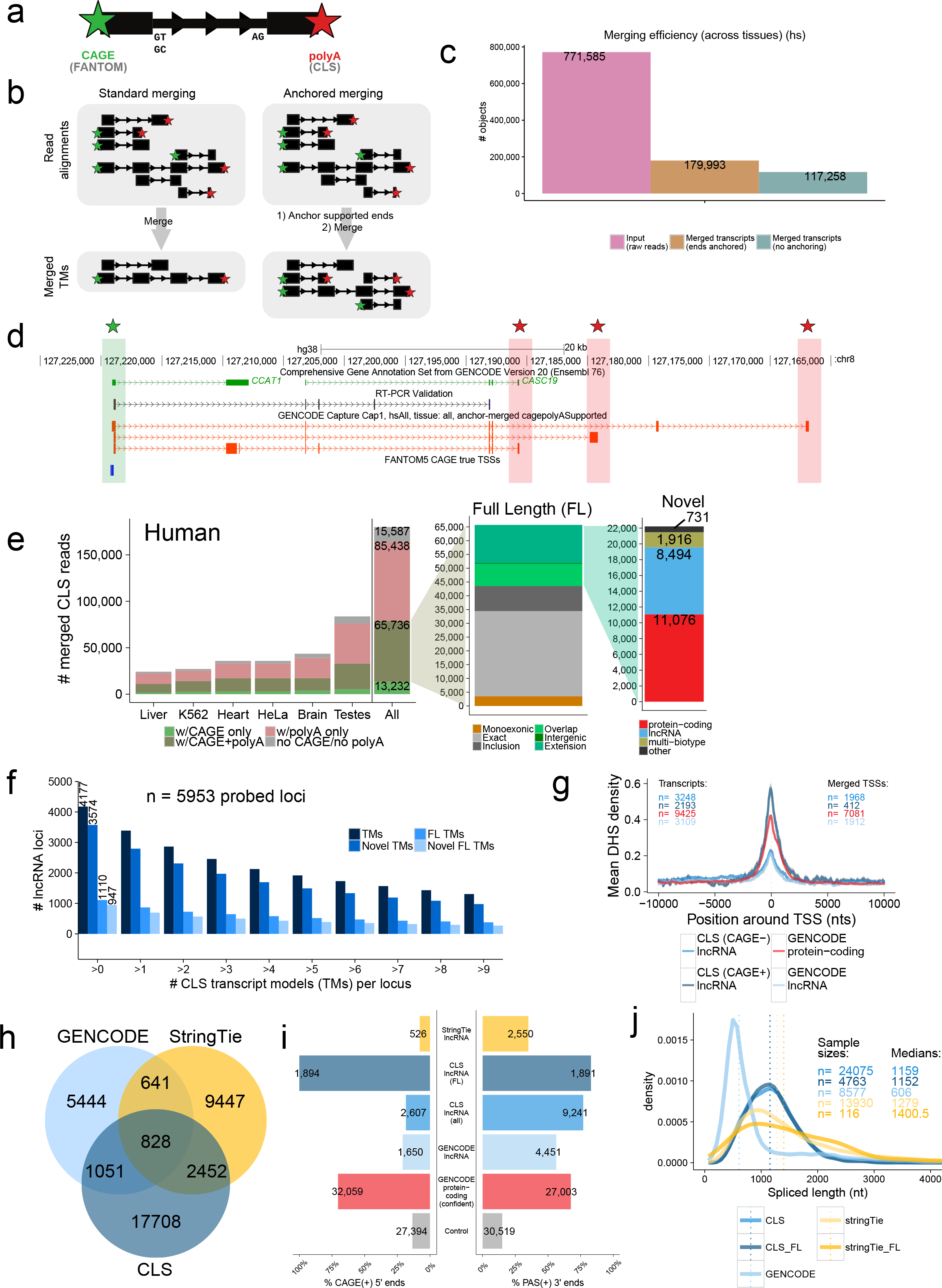
Full-length transcript annotation. **a)** 5’ and 3’ termini of transcript models (TMs) are inferred using CAGE clusters and polyA tails in ROIs, respectively. **b)** In conventional transcript merging (CM) (left), TSSs and polyA sites overlapping other exons are lost. “Anchored merging” (AM) (right) preserves such sites. **c)** AM yields more distinct TMs. *y*-axis: ROI count (pink), AM-TMs (brown), CM-TMs (turquoise). **d)** Full-length (FL) TMs at the *CCAT1* / *CASC19* locus. Red: novel FL TMs. Green/Red stars: CAGE/polyA-supported ends, respectively. An RT-PCR-amplified sequence is shown. **e)** AM-TMs for human (mouse data in Supplementary Figure 11b). *y*-axis: unique TM counts. Left: All AM-TMs, coloured by end support. Middle: FL TMs, coloured by novelty w.r.t. GENCODE. Green: novel TMs (see Methods for subcategories). Right: Novel FL TMs, coloured by biotype. **f)** Numbers of probed lncRNA loci mapped by CLS at increasing cutoffs for each category (human) (mouse data in Supplementary Figure 11c). **g)** DHS coverage of TSSs in HeLa-S3. *y*-axis: mean DHS density per TSS. Grey fringes: S.E.M. “CAGE+” / “CAGE-“: CLS TMs with / without supported 5’ ends, respectively. “GENCODE protein-coding”: TSSs of protein-coding genes. **h)** Comparing lncRNA transcript catalogues from GENCODE, CLS, and *StringTie* within captured regions. Mouse data in Supplementary Figure 12b-e. **i)** 5’/3’ transcript completeness, estimated by CAGE and upstream polyadenylation signals (PAS), respectively (human). Shown is the proportion of transcript ends with such support (“CAGE(+)”/”PAS(+)”). “Control”: random sample of internal exons. Mouse data in Supplementary Figure 12f. **j)** Spliced length distributions of transcript catalogues. Dotted line: median. Mouse data in Supplementary Figure 12c.

We developed a pipeline to merge ROIs into a non-redundant collection of transcript models (TMs). In contrast to previous approaches^4^, our “anchored merging” method preserves confirmed internal TSS or polyA sites (Figure 4b). Applying this to captured ROIs results in a greater number of unique TMs than would be identified otherwise (Figure 4c, Supplementary Figure 11a). We identified 179,993 / 129,556 transcript models across all biotypes (Supplementary Table 2), 86 / 87% of which displayed support of their entire intron chain by captured HiSeq split reads (Supplementary Table 3). In the heavily-studied *CCAT1* locus^40^, novel full-length transcripts with 5’ and 3’ support were identified (Figure 4d). CLS here suggests that adjacent *CCAT1* and *CASC19* annotations are fragments of a single gene, a conclusion supported by RT-PCR (Figure 4d).

Merged TMs can be defined by their end support: full length (“FL”, 5’ and 3’ supported), 5’ only, 3’ only, or unsupported (Figure 4b,e). We identified a total of 65,736 / 44,673 FL transcript models (Figure 4e and Supplementary Figure 11b, left panels): 47,672 (73%) / 37,244 (83%) arise from protein coding genes, and 13,071 (20%) / 5,329 (12%) from lncRNAs (Supplementary Table 2). An additional 3,742 (6%) / 1,258 (3%) represent FL models that span loci of different biotypes (listed in Figure 1b), usually including one protein-coding gene (“Multi-Biotype”). Of the remaining non-coding FL transcript models, 295 / 434 are novel, arising from unannotated gene loci. Altogether, 11,429 / 4,350 full-length structures arise from probed lncRNA loci, of which 8,494 / 3,168 (74% / 73%) are novel (Supplementary Table 2). We identified at least one FL TM for 19% / 12% of the originally-probed lncRNA annotation (Figure 4f, Supplementary Figure 11c). Independent evidence for gene promoters from DNaseI hypersensitivity sites, supported our 5’ identification strategy (Figure 4g). Human lncRNAs with mouse orthologues had significantly more FL TMs, although the reciprocal was not observed (Supplementary Figure 11d-g). This imbalance may be due to evolutionary factors (*e.g*. the appearance of novel lncRNA isoform complexity during primate evolution), or technical biases: it is noteworthy that we had access to deeper CAGE data in human than in mouse (217,516 vs 129,465 TSSs), and that human lncRNA annotations are more complete than for mouse.

In addition to probed lncRNA loci, CLS also discovered several thousand novel TMs originating from unannotated regions, mapping to probed (blue in Figure 1b) or unprobed regions (Supplementary Figures 11h-i). These TMs tended to have lower detection rates (Supplementary Figure 11j) consistent with low overall expression (Supplementary Figure 11k) and lower rates of 5’ and 3’ support than probed lncRNAs, although a small number are full length (“other” in Figure 4e and Supplementary Figure 11b, right panels).

We next compared CLS performance to the conventional, short-read CaptureSeq methodology. We took advantage of our HiSeq analysis (212/156 million reads, in human/mouse) of the same captured cDNAs, to make a fair comparison between methods. Short-read methods depend on *in-silico* transcriptome assembly: we found, using PacBio reads as a reference, that the recent *StringTie* tool outperforms *Cufflinks*, used in previous CaptureSeq projects^24,41^ (Supplementary Figure 12a). Using intron chains to compare annotations, we found that CLS identifies 69% / 114% more novel TMs than *StringTie* assembly (Figure 4h and Supplementary Figure 12b). CLS TMs are more complete at 5’ and 3’ ends than *StringTie* assemblies, and more complete at the 3’ end compared to probed GENCODE annotations (Figure 4i and Supplementary Figures 12d-h). Thus, although *StringTie* TMs are slightly longer (Figure 4j and Supplementary Figure 12c), they are far less likely to be full-length than CLS. This greater length may be due to *StringTie* producing overly long 5’ extensions, as suggested by the relatively high CAGE signal density downstream of *StringTie* TSSs (Supplementary Figure 12g-h). CLS is more sensitive in the detection of repetitive regions, identifying in human approximately 20% more repetitive nucleotides (Supplementary Figure 12i).

### Re-defining lncRNA promoter and gene characteristics

With a full-length lncRNA catalogue, we revisited the basic characteristics of lncRNA and protein-coding genes. LncRNA transcripts, as annotated, are significantly shorter and have less exons than mRNAs^5,11^. However it has remained unresolved whether this is a genuine biological trend, or simply the result of annotation incompleteness^21^. Considering FL TMs from CLS, we find that the median lncRNA transcript to be 1108 / 1067 nt, similar to mRNAs mapped by the same criteria (1240 / 1320 nt) (Figure 5a, Supplementary Figure 13a). This length difference of 11% / 19% is statistically significant (P<2×10^−16^ for human and mouse, two-sided Wilcoxon test). These measured lengths are still shorter than most annotated protein-coding transcripts (median 1,543 nt in GENCODE v20), but much larger than annotated lncRNAs (median 668 nt). There are two factors that preclude our making firm statements regarding relative lengths of lncRNAs and mRNAs: first, the upper length limitation of PacBio reads (Figure 2b); second, the fact that our size-selection protocol selects against shorter transcripts. Nevertheless we do not find evidence that lncRNAs are substantially shorter^11^. We expect that this issue will be definitively answered with future nanopore sequencing approaches.

**Figure 5:**
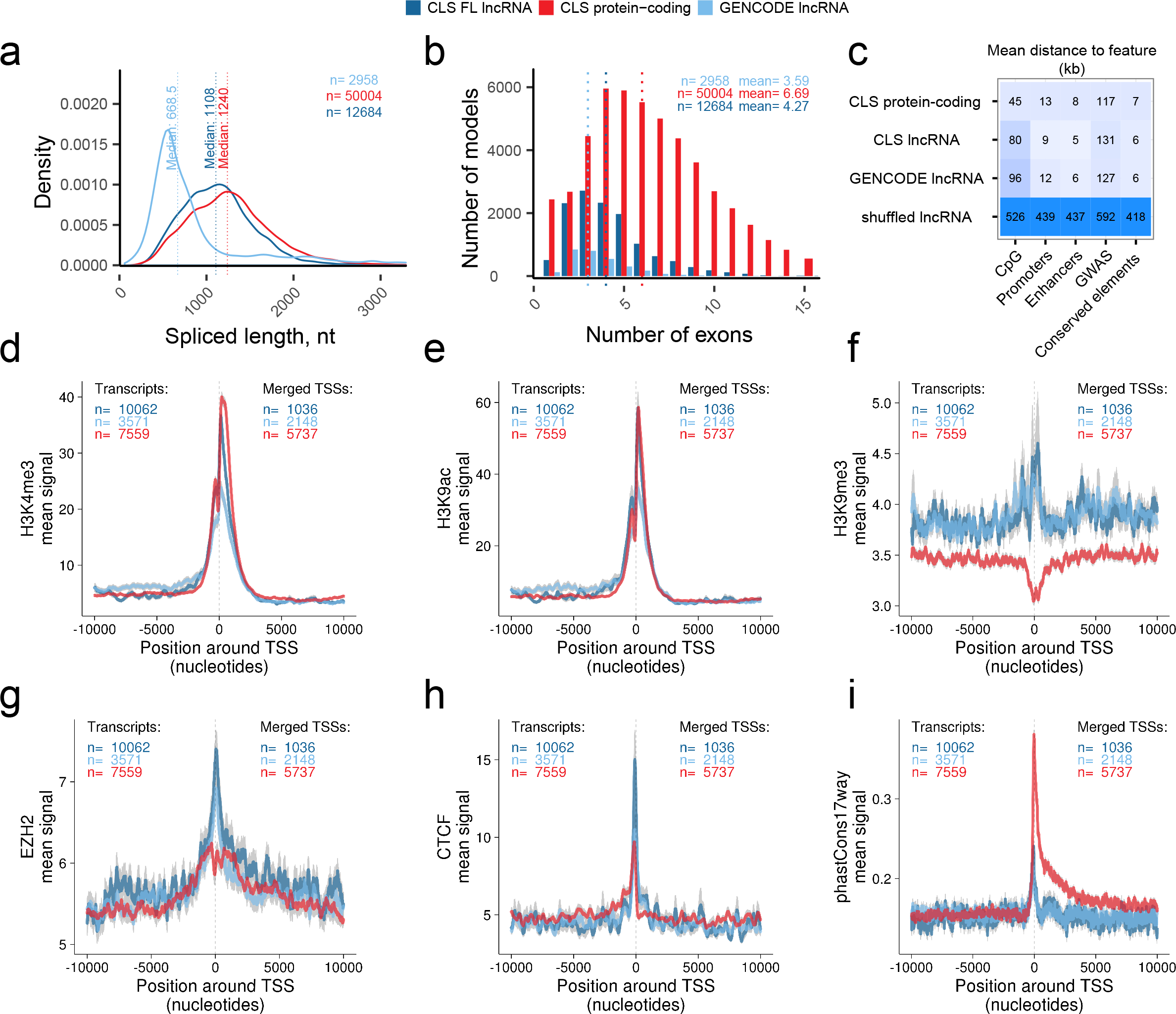
Discovery of novel lncRNA transcripts. **a)** The mature, spliced transcript length of: CLS full-length transcript models from targeted lncRNA loci (dark blue); transcript models from the targeted and detected GENCODE lncRNA loci (light blue); CLS full-length transcript models from protein-coding loci (red). **b)** The numbers of exons per full length transcript model, from the same groups as in (a). Dotted lines represent medians. **c)** Distance of annotated transcription start sites (TSS) to genomic features. Each cell displays the mean distance to nearest neighbouring feature for each TSS. TSS sets correspond to the classes from (a). “Shuffled” represent FL lincRNA TSS randomly placed throughout genome. **d) - (i)** Comparing promoter profiles across gene sets. The aggregate density of various features is shown across the TSS of indicated gene classes. Note that overlapping TSS were merged within classes, and TSSs belonging to bidirectional promoters were discarded (see Methods). The *y*-axis denotes the mean signal per TSS, and grey fringes represent the standard error of the mean. ChIP-Seq experiments are from *HeLa* cells (see Methods). *phastCons17way:* conservation scores across 17 vertebrate species. Gene sets are: Dark blue, full-length lncRNA models from CLS; Light blue, the GENCODE annotation models from which the latter were probed; Red, a subset of protein-coding genes with similar expression in *HeLa* as the CLS lncRNAs.

We previously observed a striking enrichment for two-exon genes in lncRNAs^11^. However, we have found that this is clearly an artefact arising from annotation incompleteness: the mean number of exons for lncRNAs in the FL models is 4.27, compared to 6.69 for mRNAs (Figure 5b, Supplementary Figure 13b). This difference is explained by lncRNAs’ longer exons, although they peak at approximately 150 bp, or one nucleosomal turn (Supplementary Figure 13c).

Improvements in TSS annotation are further demonstrated by the fact that FL transcripts’ TSSs are, on average, closer to expected promoter features, including promoters and enhancers predicted by genome segmentations^42^ and CpG islands, although not evolutionarily-conserved elements or phenotypic GWAS sites^43^ (Figure 5c). Accurate mapping of lncRNA promoters may provide new hypotheses for the latter’s mechanism of action. For example, improved 5’ annotation strengthens the link between GWAS SNP rs246185, correlating with QT-interval and lying in the promoter of heart- and muscle-expressed RP11-65J2 (ENSG00000262454), for which it is an expression quantitative trait locus (eQTL)^44^ (Supplementary Figure 13d-e).

Improved 5’ definition provided by CLS transcript models also allows us to compare lncRNA and mRNA promoters. Recent studies, based on the start position of gene annotations, have claimed to observe strong differences between lncRNA and mRNA promoters^45,46^. To make fair comparisons, we created an expression-matched set of mRNAs in HeLa and K562 cells, and removed bidirectional promoters. These were compared across a variety of datasets from ENCODE^12^ (Supplementary Figures 14, 15).

We observe a series of similar and divergent features of lncRNAs’ and mRNAs’ promoters. For example, activating promoter histone modifications such as H3K4me3 (Figure 5d) and H3K9ac (Figure 5e), are essentially indistinguishable between full-length lncRNAs (dark blue) and protein-coding genes (red), suggesting that, when accounting for expression differences, active promoter architecture of lncRNAs is not unique. The contrast of these findings with previous reports, suggest that the latter’s reliance on annotations alone led to inaccurate promoter identification^45,46^.

On the other hand, and as observed previously, lncRNA promoters are distinguished by elevated levels of repressive chromatin marks, such as H3K9me3 (Figure 5f) and H3K27me3^45^ (Supplementary Figures 14, 15). This may be the consequence of elevated recruitment to lncRNAs of the Polycomb Repressive Complex, as evidenced by its subunit Ezh2 (Figure 5g). Promoters of lncRNAs are also distinguished by a localised peak of insulator protein CTCF (Figure 5h). Finally, there is a clear signal of evolutionary conservation at lncRNA promoters, although lower than for protein-coding genes (Figure 5i).

Two conclusions are drawn. First, that CLS-inferred TSS have greater density of expected promoter features, compared to probed annotations, demonstrating that CLS improves TSS annotation. And second, that when adjusting for expression, lncRNA have similar activating histone modifications, but distinct repressive modifications, compared to protein-coding genes.

### Discovery of new potential open reading frames

A number of studies have suggested that lncRNA loci encode peptide sequences through unannotated open reading frames (ORFs)^47,48^. We searched for signals of protein-coding potential in FL models using two complementary methods, based on evolutionary conservation and intrinsic sequence features^49,50^ (Figure 6a, Methods, Supplementary Data 1). This analysis finds evidence for protein-coding potential in a small fraction of lncRNA FL TMs (109/1271=8.6%), with a similar number of protein-coding FL TMs displaying no evidence of encoding protein (2900/42,758=6.8%) (Figure 6b).

**Figure 6:**
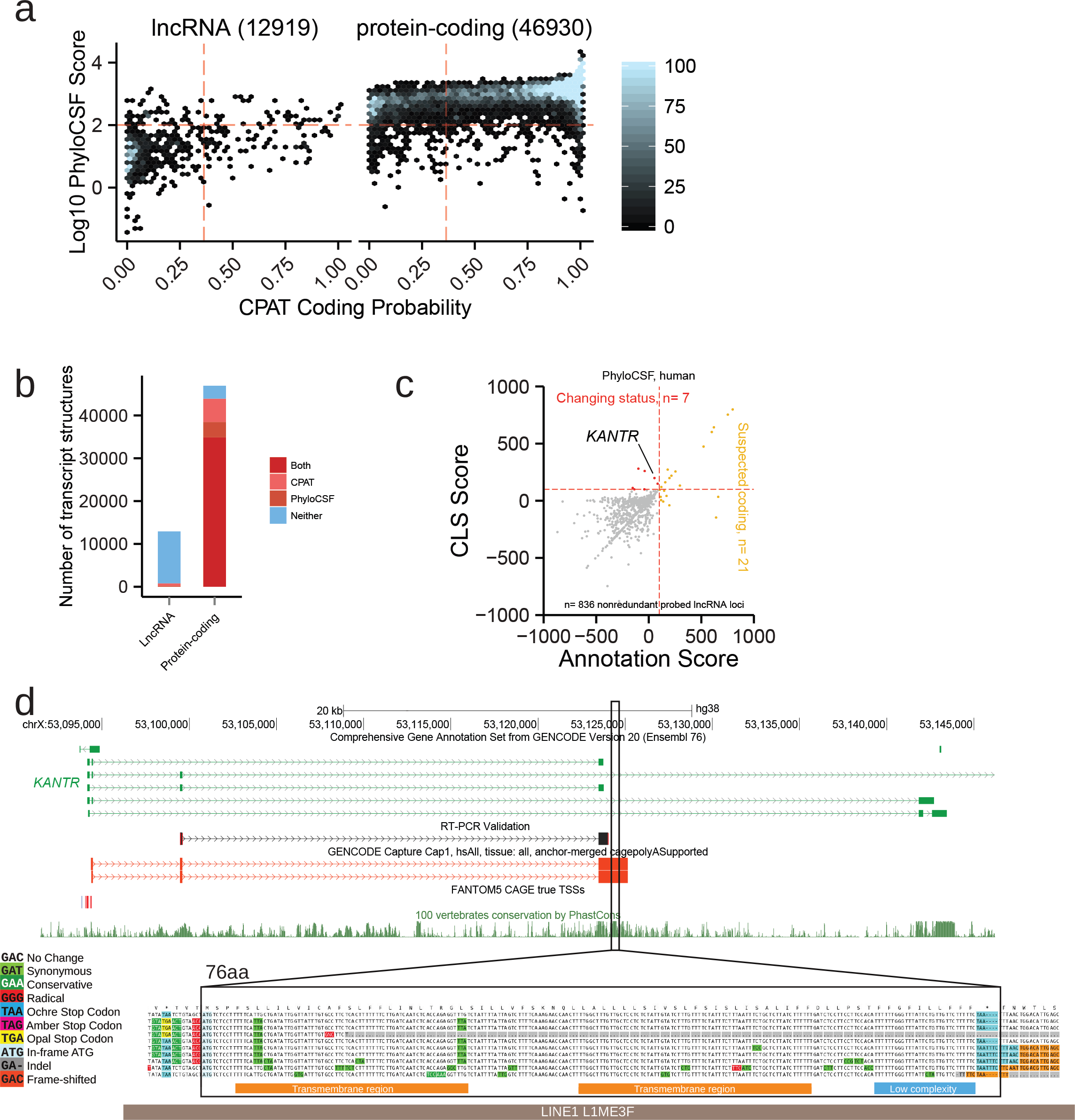
Properties of full-length lncRNAs. **a)** The predicted protein-coding potential of all full-length transcript models mapped to lncRNA (left) or protein-coding loci (right). Points represent full length (FL) transcript models (TM). *y*-axis displays the coding likelihood according to *PhyloCSF*, based on multiple genome alignments; *x*-axis displays that calculated by CPAT, an alignment-free method. Red lines indicate score thresholds, above which are considered protein-coding. TMs mapping to multiple biotypes were not considered. **b)** Numbers of classified TMs from (a). **c)** Discovery of new protein-coding transcripts in full-length CLS reads, using PhyloCSF. *x* axis: For each probed GENCODE gene annotation, score of best ORF across all transcripts; *y* axis: Score of best ORF in corresponding FL CLS TMs. Yellow: Loci from GENCODE v20 annotation predicted to encode proteins are highlighted. Red: LncRNA loci where new ORFs are discovered as a result of CLS transcript models. **d)** *KANTR*, example of an annotated lncRNA locus novel protein-coding sequence is discovered. The upper panel shows the structure of the lncRNA and the associated ORF (highlighted region) falling within novel FL CLS transcripts (red). Note how this ORF lies outside existing annotation (green), and overlaps a highly-conserved region (see PhastCons conservation track, below). Shown is a sequence obtained by RT-PCR (black). The lower panel, generated by *CodAlignView* (see URLs), reveals conservative substitutions in the predicted 76 aa ORF consistent with a functional peptide. High-confidence predicted SMART^53^ domains are shown as coloured bars below. This ORF lies within and antisense to a L1 transposable element (grey bar).

CLS FL models support a reclassification of protein-coding potential for five distinct gene loci (Figure 6c, Supplementary Figure 16a, Supplementary Data 2). A good example is the *KANTR* locus, where extension by CLS (supported by independent RT-PCR) identifies a placental mammal-conserved 76aa ORF with no detectable protein orthologue^51^. It is composed of two sequential transmembrane domains (Figure 6d, Supplementary Figure 16e), and derives from a LINE1 transposable element. Another case is *LINC01138*, linked with prostate cancer, where a potential 42 aa ORF is found in the extended transcript^52^. We could not find peptide evidence for translation of either ORF (see Methods). Whole-cell expression, as well as cytoplasmic-to-nuclear distributions, also showed that potentially protein-coding lncRNAs’ behaviour is consistently more similar to annotated lncRNAs than to mRNAs (Supplementary Figures 16b-d). Hence, CLS will be useful in improving biotype annotation of the small minority of lncRNAs that may encode proteins.

## Discussion

We have introduced an annotation methodology that resolves the competing needs of quality and throughput. Capture Long Read Sequencing produces transcript models with quality approaching that of human annotators, yet with throughput comparable to *in-silico* transcriptome reconstruction. CLS improves upon existing assembly-based methods not only due to confident exon connectivity, but also through (1) far higher rates of 5’ and 3’ completeness, and (2) carrying encoded polyA tails.

In economic terms, CLS is also competitive. Using conservative estimates, with 2016 prices ($2460 for 1 lane of PE125bp HiSeq, $500 for 1 SMRT), and including the cost of sequencing alone, we estimate that CLS yielded one novel, full-length lncRNA structure for every $8 spent, compared to $27 for conventional CaptureSeq. This difference is due to the greater rate of full-length transcript discovery by CLS.

Despite its advantages, CLS remains to be optimised in several respects. First, the capture efficiency for long cDNAs can be improved by several-fold. Second, various technical factors limit the completeness of CLS TMs, including: sequencing reads that remain shorter than many transcripts; incomplete reverse transcription of the RNA template; degradation of RNA molecules before reverse transcription. Resolving these issues will be important objectives of future protocol improvements, and only then can we make definitive judgements about lncRNA transcript properties. In recent, unpublished work we have further optimised the capture protocol, pushing on-target rates to around 35% (see details in Methods). However the most dramatic gains in cost-effectiveness and completeness of CLS will be made by advances in sequencing technology. The latest nanopore cDNA sequencing promises to be ~150-fold cheaper per read than PacBio technology (0.01 vs 15 cents/read, respectively).

Full-length annotations have provided the most confident view to date of lncRNA gene properties. These are more similar to mRNAs than previously thought, in terms of spliced length and exon count^11^. A similar trend is seen for promoters: when lncRNA promoters are accurately mapped by CLS and compared to expression-matched protein-coding genes, we find them to be surprisingly similar for activating modifications. This suggests that previous studies, which placed confidence in annotations of TSS, should be reassessed^45,46^. On the other hand lncRNA promoters do have unique properties, including elevated levels of repressive histone modification, recruitment of Polycomb group proteins, and interaction with the insulator protein CTCF. To our knowledge, this is the first report to suggest a relationship between lncRNAs and insulator elements. Overall, these results suggest that lncRNA gene features *per se* are generally comparable to mRNAs, after normalising for their differences in expression. Finally, extended TMs do not yield evidence for widespread protein-coding capacity encoded in lncRNAs.

Despite success in mapping novel structure in annotated lncRNAs, we observed surprisingly low numbers of transcript models originating in the relatively fewer numbers of unannotated loci that we probed, including ultraconserved elements and developmental enhancers. This would suggest that, at least in the tissue samples probed here, such elements do not give rise to substantial numbers of lncRNA-like, polyadenylated transcripts.

In summary, by resolving a longstanding roadblock in lncRNA transcript annotation, the CLS approach promises to accelerate progress towards an eventual “complete” mammalian transcriptome annotation. These updated lncRNA catalogues represent a valuable resource to the genomic and biomedical communities, and address fundamental issues of lncRNA biology.

## URLs

CLS data portal: https://public_docs.crg.es/rguigo/CLS/.

Pre-loaded CLS UCSC Genome Browser track hub: http://genome-euro.ucsc.edu/cgi-bin/hgTracks?hubUrl=http://public_docs.crg.es/rguigo/CLS/data/trackHub//hub.txt *CodAlignView:* https://data.broadinstitute.org/compbio1/cav.php.

ENCODE mycoplasma contamination guidelines:https://www.encodeproject.org/documents/60b6b535-870f-436b-8943-a7e5787358eb/@@download/attachment/Cell_Culture_Guidelines.pdf

## Acknowledgements

We thank members of the Guigó laboratory for their valuable input and help when handling samples, analysing data and writing the manuscript, including Emilio Palumbo, Ferran Reverter, Alessandra Breschi, Dmitri Pervouchine, Carme Arnan and Francisco Camara. We wish to thank Lluis Armengol (qGenomics) for advice on RNA capture, Diego Garrido (CRG) for help with eQTL analysis, Sarah Bonnin (CRG) for help with data manipulation in R, Irwin Jungreis (MIT) for advice on PhyloCSF. James Wright and Jyoti Choudhary (Sanger Institute) helped in searching for peptide hits to putative coding regions. Sarah Djebali (INRA, France) kindly made available the *Compmerge* utility. This work and publication were supported by the National Human Genome Research Institute of the National Institutes of Health (grant numbers U41HG007234, U41HG007000 and U54HG007004) and the Wellcome Trust (grant number WT098051). RJ was supported by Ramón y Cajal RYC-2011-08851. Work in laboratory of RG was supported by Awards Number U54HG0070, R01MH101814 and U41HG007234 from the National Human Genome Research Institute. This research was partly supported by the NCCR RNA & Disease funded by the Swiss National Science Foundation (to RJ). We thank Romina Garrido (CRG) for administrative support. We acknowledge support of the Spanish Ministry of Economy and Competitiveness, ‘Centro de Excelencia Severo Ochoa 2013-2017’ (SEV-2012-0208) and the support of the CERCA Programme, Generalitat de Catalunya.

## Author contributions

RJ, RG, JH, AF, BU-R and JL designed the experiment. SC generated cDNA libraries and performed the Capture. CD and TRG performed the PacBio sequencing of Capture libraries. JL and BU-R analysed the data under the supervision of RG and RJ. RJ wrote the manuscript, with contributions from JL, BU-R and RG. SP-L and AA performed the RT-PCR experiments.

## Online Methods

### Abbreviations

FL: full length
HCGM: High-Confidence Genome Mapping
ROI: read of insert, *i.e.* PacBio reads
SJ: splice junction
SS: splice site
TM: transcript model
TSS: Transcription Start Site
UMD-ROI: Uniquely Mapped and Demultiplexed ROI

## Library Design

### Design of human capture probes

All designs were based on the GENCODE^10^ version 20 annotation in human genome build hg38. For probe design, a target annotation was prepared in FASTA format and composed of the following sets of features. In each case, the entire set of feature of each class was taken as a starting point, unless otherwise stated, and where necessary were lifted over to the hg38 assembly. Features overlapping protein coding gene loci were removed. Intergenic lncRNAs were extracted from the GENCODE v20 annotation, being all those genes having no single transcript that overlaps or lies within 5 kb of any protein-coding gene. For small RNA loci, a 1 kb window centered on the small RNA was targeted.

At this stage, the expression of candidate regions was quantified using HBM/ENCODE RNA-Seq data from appropriate human tissues and cell lines. We noticed that the top 20 most expressed features (mean expression across samples) produced approximately 71% of sequencing reads (Supplementary Figure 17) – these were removed, in order to favour rarer transcripts. A number of controls were added to the design. 100 protein coding genes, with steady-state levels matched to the distribution of lncRNAs, were included. 100 random genomic regions of 1 kb from the *E. coli* genome were included. 100 intergenic regions of 1 kb each with no evidence from ENCODE ChromHMM for any transcriptional or regulatory activity^54^. Finally, out of the 92 ERCC sequences, we removed the top 8 most concentrated, and selected half (42) of the remainder such that they evenly covered the concentration distribution. In total the design targeted 14,667 regions, which corresponded to ~15.5 Mb of human genome (hg38) and exons of 9,560 lincRNAs from 5,953 loci. The summary information for selected transcript targets in human is provided in Supplementary Table 4. Statistics on probed gene loci are presented in Figure 1b.

All targets were combined into a single FASTA file and submitted to Roche NimbleGen (Madison, WI) for probe design. The oligonucleotide-probes were designed and synthesised as a SeqCap EZ Choice XL Library according to manufacturer’s protocol. The oligonucleotide probes covered 86.6% of target regions directly, with an estimated 96.1% of target regions successfully targeted. Roche Nimblegen’s policy prohibited release of SeqCap’s probe coordinates, but the design is available on request.

### Design of mouse capture probes

Mouse library design was carried out essentially as for human with the following differences. All designs were based on the GENCODE version M3 annotation in genome build mm10. Candidate lncRNAs were filtered to remove those overlapping any protein coding gene within 5kb. Homology-based predictions of mouse orthologues of human lncRNA were obtained using the PipeR pipeline^29^. As before, the top 20 most expressed lncRNAs, as estimated using RNAseq data from ENCODE^31^ were removed. The final design covered 8,708 regions, including 2,817 GENCODE vM3 lincRNA transcripts from 1,920 loci. The covered regions corresponded to 8.3 Mb. The summary information for selected transcript targets in mouse is provided in Supplementary Table 5. Statistics on probed gene loci are presented in Figure 1b.

Designed oligonucleotide probes covered 76.3% of target regions directly and 85.0 % of target regions successfully targeted. Oligonucleotide-probes were synthesised as a Roche NimbleGen SeqCap EZ Choice XL Library. Roche Nimblegen’s policy prohibited release of SeqCap’s probe coordinates, but the design is available on request.

## Sample Preparation

### RNA samples

Commercial total RNA samples were obtained for 4 different adult human (Ambion AM6000) and mouse (Clontech 636644) tissues: heart, testes, liver and brain. From the same panel also came mouse E7 and E15 samples. Human K562 and HeLa RNA was obtained directly from members of the ENCODE consortium^31^. Neither cell line used in this paper is listed in the database of commonly misidentified cell lines maintained by ICLAC. Cell lines were not authenticated. Cell lines were tested for mycoplasma contamination as per ENCODE guidelines (see URLs). The integrity of samples were tested by Bionanalyzer (Agilent) and all had values of 8.5 or higher. To 4 μg of each RNA sample, we added 4 μl of 1:100 diluted ERCC mix (Ambion 4456740) according to manufacturer’s protocol (Supplementary Table 6). Mix 1&2 were assigned to samples as shown below. The samples containing ERCC controls were ribodepleted with Ribo-Zero (Epicentre MRZE724), and successful rRNA removal was validated by Bioanalyzer.

### cDNA synthesis

Full-length cDNA was synthetized via reverse-transcription of ribosome-depleted RNA samples using SMARTer^®^ PCR cDNA Synthesis Kit (Clontech 634926) and Advantage^®^2 PCR Kit (Clontech 639206). Each cDNA was synthetized using 3.5 μl of ribosome-depleted RNA according to manufacturer protocol, performing two independent cDNA synthesis reactions for each sample. It should be noted that cDNA was primed using oligodT. The adapter used in the cDNA library construction sequences (“SMART IV Oligonucleotide” and “CDS III/3’ PCR Primer”) are to be found in Supplementary Data 3.

All first strand RNA obtained from the reaction was used for second strand synthesis. The synthesis cycling protocol was modified from that specified by the manufacturer, increasing the extension time from 3 to 6 minutes to favour the synthesis of long strands. After protocol optimization, a total of 18 cycles were used to obtain the full-length cDNA libraries. Resulting cDNA was quantified using NanoDrop^®^ ND-1000 full-spectrum spectrophotometer (Thermo Scientific). The library length and quality was also verified by Bioanalyzer.

## Capture

### Library Preparation

cDNA samples were used to create barcoded, full length libraries. The two aliquots of cDNA obtained in the preceding step were pooled, of which 1 ug was used for library preparation. One adenine was added to blunt cDNA 3’ extremities and Illumina Truseq adapters were ligated. Different barcoded adaptor hexamer indexes were used to discriminate each sample (Supplementary Table 7 and Supplementary Data 3). The overall structure of cDNA libraries is represented schematically in Supplementary Figure 2c.

The library was amplified for 10 PCR cycles modifying standard Kapa Biosystems PCR conditions (Low throughput Library prep - Kapa Biosystems KK8232). The PCR extension step was increased to 3 min to allow long fragments to be fully amplified. The quality and the length of libraries was checked using Agilent 2100 Bioanalyzer. Library quantification was performed using Qubit^®^ dsDNA BR assays (ThermoFisher). For each cDNA sample, an additional Covaris-fragmented, Illumina sequencing library was prepared for MiSeq and HiSeq sequencing according to standard protocols.

Standard Illumina 6mer indexes were used here, to be compatible with blocking oligonucleotides in the SeqCap capture protocol (see below). It should be pointed out that the use of these relatively short indexes led to loss of information during later demultiplexing steps. Improving this issue by using standard 16-nt PacBio indexes should be a priority in future versions of CLS.

### Sample pooling

Samples were pooled separately by species, such that all 6 human libraries were mixed at equimolar ratios, and similarly for mouse. The final amount of each pool was one microgram.

### cDNA capture

Human and mouse pools were dried and prepared for hybridisation to NimbleGen SeqCap EZ Choice XL Library capture probes, according to manufacturer’s protocol (SeqCap EZ Library SR User’s Guide - Version 5.0). Hybridisation was carried out for 72 hours. Altogether 5 separate, parallel captures were performed for each species: 4 were used for subsequent PacBio sequencing, with one remaining sample used for Illumina sequencing.

Subsequent to the presented work, we have managed to further optimise the efficiency of this capture process, by implementing four changes to the described protocol:

1. Drying cDNA for resuspension prior to capture: temperature at 60°C instead of 55°C.
2. Hybridisation incubation time: 20h instead of 72h.
3. Washing steps after capture: use of water bath instead of dry bath.
4. Blockers: Additional blockers targeting the SMARTer adaptors used during library construction (sequences in Supplementary Data 3, “SMARTer_blocker” and “SMARTer_5p_PCR_blocker”).

### Amplification and quality control of captured cDNA

Following hybridisation, human and house pools were washed to eliminate nonspecific hybridization using m-280 streptavidin Dynabeads (Invitrogen 11205D) following recommendations of Roche protocol. Human and Mouse washed pools were PCR amplified using KAPA HotStart ReadyMix- 2X (KapaBiosysthems KK1006). Two independent PCR reactions containing half of washed pool each were prepared to avoid PCR duplicates. 18 cycles of PCR were performed, with an increased extension step of 3 minutes to allow long fragments to be fully amplified. The length of post-capture PacBio and Illumina libraries was verified by Bioanalyzer, and quantity by Qubit.

## Pacific Biosciences (PacBio) Sequencing of Captured cDNA

### Pooling

After quantification and quality control the 4 post-capture libraries were pooled together by species to obtain one unique human and one unique mouse pool. The 110 μl of each sample were again quantified by Qubit^®^ dsDNA BR assays (ThermoFisher): 12.3 μg for human and 9.57 μg for mouse.

### Size Selection

Samples were subsequently size-selected using E-gel (Invitrogen) to obtain 3 different ranges: 1000-1500 bp, 1500-2500 bp and >2500 bp. Two shorter fractions of 200-500 bp, 500-1000 bp were collected, but following preliminary sequencing data, it was decided not to scale them up, due to the large number of reads in this size range obtained in the larger fractions. Following size selection, each size fraction was dried and resuspended with 20 μl of water and quantified by Qubit^®^ dsDNA BR assays (ThermoFisher). These samples were then amplified again by PCR (4 cycles) using Kapa HiFi HotStart (kapaBiosystems) in order to reach the required amount for PacBio library preparation. The quality and the length of obtained libraries was verified using Bioanalyzer and Qubit.

The efficiency of size selection was also checked by analysis of spike-in sequences (Supplementary Figure 1d). For each size-selected captured library, and for pre-capture libraries, the sequencing efficiency was calculated as a function of spike-in sequence length. Sequencing efficiency was defined for each spike-in sequence as: (Number of reads) / (molar concentration * sequence length * total read count). This showed that, as expected, size-selection boosted the sequencing of longer templates.

### PacBio library preparation

Approximately 2μg of each of the size-fractionated and amplified DNA was used for each of human and mouse pools, for a total of 6 (3×2) distinct samples. Sizes and concentrations were verified by Bioanalyzer. PacBio libraries were constructed for each sample using Kit #100-250-100 (Pacific Biosciences Inc.) as per the manufacturer’s protocol. Briefly, this involves polishing the PCR amplicon ends to ‘blunt’ them, ligating the SMRTbell adapters, removing linear (non-ligated) fragments of DNA, AMPure bead purification, followed by Bioanalyzer analysis to assess the size distribution and Qubit quantifications.

### PacBio sequencing and collection of Post-Capture data

The PacBio libraries were each run on an initial SMRTcell to assess their respective performance and optimal sequencing concentration. Those that performed well were then scaled-up to an additional 20 SMRTcells for deep data collection. The PacBio reagents and metrics used for each sample are listed in Supplementary Table 8. The sequencing was performed on a PacBio RSII instrument. Upon completion of the sequencing, SMRTcells from a given library were aggregated on the SMRTportal and the PacBio post-processing method “RS_ReadsOfInsert.1” were run on each aggregated sample to obtain Reads of Inserts (ROIs) for downstream processing. This yielded a single FASTQ file per library.

### HiSeq sequencing of Captured cDNA

Post-capture Illumina cDNA libraries were sequenced on a HiSeq 2500 machine (2×125 nucleotides, v4, high output mode). One sequencing lane was generated per species at a depth of ~212M (human) and ~156M (mouse) pairs of reads. Read pairs were demultiplexed using Illumina software. Note that these libraries were unstranded, and Covaris-fragmented before capture.

### Demultiplexing of ROIs according to sample barcodes

As previously mentioned, PacBio reads contained Illumina Truseq adapters: universal (59 nt) and indexed (65 nt) that flank targeted cDNAs (Supplementary Figure 2c). To demultiplex samples (i.e., determine the tissue of origin of each ROI), for each adapter its middle 26-nt was selected. Each of the 26-mers derived from the indexed adapters contained the hexamer barcode in the center. GEM mapper^55^ was employed to demultiplex samples. PacBio reads were compiled into a FASTA file (one file per species) and indexed by GEM. Mapping the middle 26-mer of indexed adapters to the PacBio read allowed us to assign it its tissue origin. The additional presence of the universal adapter within ROIs was used to confirm the completeness of the insert. The GEM-based demultiplexing procedure allowed up to 3 mismatches (`−m 0.1`) and 3 indels (`−e 0.1`), accurate identification of the barcodes. Following non-default GEM parameters were used during the mapping step: `−T 3 --max-big-indel-length 0 −s 3 −D 4`. We filtered out “chimeric” ROIs (that is, reads arising from the concatenation of inserts during adapter ligation) by removing those reads that contained more than one indexed or more than one Universal TruSeq Illumina adapter sequence.

Overall, we could demultiplex 1,627,322 and 1,509,374 ROIs in human and mouse, respectively (Figure 2a and Supplementary Figure 2b). As presented in Supplementary Figure 2d, only a minute fraction of human ROIs were assigned a mouse barcode (and reciprocally), underlining the high specificity of the demultiplexing procedure.

### Read mapping

All read-to-genome alignments were performed on genome assemblies GRCh38/hg38 (human) and GRCm38/mm10 (mouse). Mapping of ROIs from post-capture PacBio libraries to human and mouse genomes (in addition to sequences of 96 ERCC Spike-In Controls) was carried out using STAR^56^ (v.2.4.0.1) compiled for long reads. To improve splice junction mapping accuracy, a reference annotation was provided as a guide to the aligner. Reference annotation for human was built using GENCODE v20 set and sequences of all other targeted regions. For mouse, exonic sequences of *PipeR* predictions along with sequences of all other additional targets were added to the reference annotation of GENCODE vM3. The following non-default parameters where used during the mapping step: --outFilterMultimapScoreRange 20 --outFilterScoreMinOverLread 0 --outFilterMatchNminOverLread 0.5 --outFilterMismatchNmax 1000 --winAnchorMultimapNmax 200 --seedSearchStartLmax 50 --seedPerReadNmax 100000 --seedPerWindowNmax 100 --alignTranscriptsPerReadNmax 100000-alignTranscriptsPerWindowNmax --genomeSAsparseD 4 -- outSAMunmapped Within --runThreadN 6.

For analysis of MiSeq (pre-capture cDNA) and HiSeq (post-capture) data, FASTQ files were aligned to the human and mouse genomes (plus the sequences of 96 ERCC Spike-In Controls), using STAR^56^ (v.2.4.0.1) compiled for short reads. Again the reference annotations described above were employed to guide the mapper. In order to maximize the mapping rate, the mates of each pair of reads were aligned separately. The following non-default STAR parameters were specified: `--outFilterMismatchNoverLmax 0.04 --alignIntronMin 20 -- alignIntronMax 1000000 --alignMatesGapMax 1000000 -- outSAMunmapped Within --runThreadN 6`. HiSeq mapping statistics are summarized in Supplementary Figure 2l.

## Analysis of CLS performance, on-target enrichment

### RNA capture on-target enrichment

The overall RNA capture performance was evaluated by calculating an on-target rate in both MiSeq pre- and PacBio post-capture libraries. The on-target rate was defined as the ratio of the number of distinct ROIs mapping into targeted genomic regions (excluding ERCC RNA spike-in controls) over the total number of mapped ROIs. The number of reads overlapping targeted regions was calculated directly from the STAR BAM file using `bedtools intersect`^57^. Overlap was defined as ≥1 bp of intersection between the sequencing read and the exonic span of a feature, on the same strand. The overall on-target fold enrichment was computed as the on-target rate in the post-capture library, divided by the on-target rate in the pre-capture library.

Enrichment was calculated separately by reference to two distinct sequencing datasets of post-capture cDNA: (a) the main PacBio reads; (b) Illumina MiSeq of the same material. Figure 2d shows data for enrichments calculated using the latter data: MiSeq post-capture vs MiSeq pre-capture. Equivalent enrichments for the former comparison (PacBio post-capture vs MiSeq pre-capture) are: 16.6- / 11.1-fold, for human/mouse.

We compared CLS enrichments to a previous Capture Short-read Sequencing (CSS) study^24^. We focused our analysis on the CSS tissues that were also assayed in CLS (that is, human Brain, Heart, Liver and Testis), and computed on-target rates on lincRNAs more than 5kb away from any protein-coding gene in both studies, based on GENCODE v20 and v19 for CLS and CSS, respectively. CSS pre-capture rates were estimated using pre-capture MiSeq libraries generated in the present work, and remapped to hg19 / GENCODE 19. Across the 4 tissues studied, CLS outperformed CSS both in terms of on-target enrichment (in all samples) and post-capture on-target rate (in Brain and Testis only) (Supplementary Figure 2f-g).

### Breakdown of sequencing reads by gene biotype

Both the human and mouse genomes, as well as ERCC spike-in sequences, were segmented into distinct classes of locus regions, according to their gene biotype annotation and capture status (*i.e*., On-target vs Off-target). The on- and off-target categories correspond to standard, GENCODE-annotated gene biotypes (in simplified categories, as described in the Supplementary Note, in addition to “Other”, which comprises mitochondrial genes), while the “Intergenic” class includes all non-targeted and unannotated genome segments. Next, we calculated the proportion of pre- and post-capture MiSeq reads originating from each genome partition using the read BAM files and the `bedtools coverage` utility^57^. Note that when a given read overlapped multiple regions of distinct biotype classes, it was counted in each of these classes separately. Secondary targets (*i.e*., genes not targeted *per se*, yet overlapping targeted regions) were included in on-target biotype sub-classes. The following additional hierarchical rules were applied in the assignment: the highest priority in the read classification was given to capture-targeted (“On-target”), then “Off-target”, and finally “Intergenic” classes; these three categories are mutually exclusive.

### Comparison of capture protocols and long cDNA capture efficiency

We wished to compare the performance of the CLS protocol to other methods. Performance was judged by (1) the % of reads in post-capture cDNA originating from a targeted region (“on-target” rate), and (2) the enrichment, defined as the ratio of on-target rates in post/pre capture cDNA. In all experiments, the off-the-shelf SeqCap RNA lncRNA Enrichment Kit (Roche) was used. Four distinct experiments were performed. For each one, the same aliquot of human kidney total RNA was used, and sequencing was performed using Illumina MiSeq. The experiments were:

1. Original CLS protocol (as used and described here), PolyA-selected, unfragmented.
2. Improved CLS protocol, PolyA-selected, unfragmented.
3. Improved CLS protocol, Total RNA, unfragmented.
4. Roche SeqCap RNA protocol, Total RNA, fragmented.

“Improved” CLS incorporated several adjustments designed to boost enrichment: use of Lo-bind tubes, drying step at 60°C, shorter incubation time, use of Smarter blockers, use of water bath at 47°C for post capture washes.

Findings are presented in Supplementary Figure 2h-i, and together suggest that capturing long cDNAs yields lower on-target efficiency.

## Supplementary Methods

Available in the Supplementary Note.

### Code availability

All computer code used in this study is available from the authors upon request. Most programs are deposited on GitHub, as specified in the relevant sections of the text.

### Data availability

Raw and processed data is deposited in the Gene Expression Omnibus under accession GSE93848. RT-PCR validation sequences are available as Supplementary Data 4. Genome-aligned data were assembled into a public Track Hub, which can be loaded into the UCSC Genome Browser (see URLs).

